# Context-specific inhibition of mitochondrial ribosomes by phenicol and oxazolidinone antibiotics

**DOI:** 10.1101/2024.08.21.609012

**Authors:** Brianna Bibel, Tushar Raskar, Mary Couvillion, Muhoon Lee, Jordan I. Kleinman, Nono Takeuchi-Tomita, L. Stirling Churchman, James S. Fraser, Danica Galonić Fujimori

## Abstract

The antibiotics chloramphenicol (CHL) and oxazolidinones including linezolid (LZD) are known to inhibit mitochondrial translation. This can result in serious, potentially deadly, side effects when used therapeutically. Although the mechanism by which CHL and LZD inhibit bacterial ribosomes has been elucidated in detail, their mechanism of action against mitochondrial ribosomes has yet to be explored. CHL and oxazolidinones bind to the ribosomal peptidyl transfer center (PTC) of the bacterial ribosome and prevent incorporation of incoming amino acids under specific sequence contexts, causing ribosomes to stall only at certain sequences. Through mitoribosome profiling, we show that inhibition of mitochondrial ribosomes is similarly context-specific – CHL and LZD lead to mitoribosome stalling primarily when there is an alanine, serine, or threonine in the penultimate position of the nascent peptide chain. We further validate context-specific stalling through in vitro translation assays. A high resolution cryo-EM structure of LZD bound to the PTC of the human mitoribosome shows extensive similarity to the mode of bacterial inhibition and also suggests potential avenues for altering selectivity. Our findings could help inform the rational development of future, less mitotoxic, antibiotics, which are critically needed in the current era of increasing antimicrobial resistance.

## INTRODUCTION

Bacteria are acquiring resistance to mainstay antibiotics at an alarming rate, necessitating the development of novel compounds^1^. Oxazolidinones, including the FDA-approved linezolid (Zyvox) and tedizolid (Sivextro), are a recently-introduced class of synthetic antibiotics that show activity against multiple drug-resistant pathogens. Unfortunately, like several other antibiotics designed to target bacterial ribosomes, including the phenicol antibiotic chloramphenicol, oxazolidinones can cause mitochondrial toxicity thought to be due to off-target inhibition of mitochondrial ribosomes (mitoribosomes)^2^. Mitoribosomes translate 13 proteins (from 11 transcripts), all of which are crucial components of the mitochondrial electron transport chain. Inhibition of mitochondrial translation has thus been implicated in lactic acidosis, myelosuppression, peripheral neuropathy, and other serious, sometimes life-threatening, complications^3–5^. These restrict the use of oxazolidinones and, along with growing resistance, fuel an active search for less mitotoxic alternatives. Mitoribosomes are susceptible to phenicol- and oxazolidinone-mediated inhibition due to their structural similarity to bacterial ribosomes^6^; however, the extent to which the mechanisms of binding and inhibition are conserved is unknown.

Structural and biochemical studies have elucidated the basis of bacterial ribosomal inhibition by phenicols and oxazolidinones^7–10^. Oxazolidinones and phenicols bind to the A site of bacterial ribosomes’ peptidyl transfer center (PTC)^11^. These antibiotics were previously thought to inhibit translation globally by preventing accommodation of the A site tRNA^8^; however, it was recently discovered that chloramphenicol (CHL) and linezolid (LZD) instead act as context-dependent inhibitors of bacterial translation. More specifically, LZD and CHL cause preferential stalling during the translation of specific nascent peptides, chiefly when an alanine (Ala), and to a lesser extent serine (Ser) or threonine (Thr), residue is in the penultimate position of the nascent chain (−1 with respect to the P site, or P-1)^7^. We and others recently showed that this context-dependency is due to steric constraints of the antibiotic-bound ribosome and stabilizing CH-π interaction between the alanine’s methyl group and the aryl ring of the oxazolidinone or phenicol^9,10^. Analogous interactions via Ser and Thr’s side chains similarly stabilize antibiotic binding in the bacterial ribosome^9^, leading to the secondary Ser/Thr bias, which is more pronounced for CHL than LZD^7^.

Due to the endosymbiotic origins of mitochondria, the PTC is well-conserved between bacterial and mitochondrial ribosomes^12^. Accordingly, mitoribosomes are susceptible to inhibition by CHL, LZD, and other oxazolidinones^13^. The ability of CHL to inhibit mitochondrial translation has long been known – it has led to its broad disuse as an oral antibiotic, in parallel with its broad use as an experimental tool to block mitochondrial translation in a laboratory setting. Multiple studies have found that oxazolidinones inhibit mitochondrial translation as well^13,14^. Despite these functional data, no structures of phenicol or oxazolidinone-bound mitoribosomes are available, although crosslinking data has identified a binding site of linezolid on the mitoribosome analogous to that of the bacterial ribosome^6^. Thus, the precise binding mode of CHL and oxazolidinones to mitoribosomes remains unknown. Furthermore, it is yet to be elucidated if these antibiotics act as global inhibitors of translation or whether inhibition happens in a nascent peptide-dependent manner at the mitoribosome.

Here, using a combination of mitoribosome profiling, in vitro translation assays, and cryo-electron microscopy (cryo-EM), we characterize the mechanism by which chloramphenicol and linezolid inhibit mitochondrial ribosomes. Through mitoribosome profiling in human cells, we discover that CHL and LZD do indeed inhibit mitoribosomal translation in a context-dependent manner. We find that, in the presence of these antibiotics, mitoribosomes stall mainly on sequences where there is an alanine (or to a lesser extent serine or threonine) in the penultimate position of the nascent chain, similar to the preference seen in bacterial translation inhibition. We confirm the differential and sequence-dependent nature of such stalling in a fully reconstituted mammalian mitochondrial translation system. Finally, to investigate a potential structural basis for such inhibition, we solve the structure of the large (39S) subunit of a human mitoribosome bound to LZD. This structure shows a binding site and binding interactions that are largely well-conserved to those of bacterial ribosomes. However, it also reveals an altered linezolid-C5 tail conformation and an extended water network that is absent in bacterial ribosomes. Together, these results imply that, while the structural basis for the inhibition of mammalian mitoribosomes by LZD is similar to that of bacterial ribosomes, distinct features of its binding to mitoribosomes suggest the potential for the design of next-generation oxazolidinone antibiotics with lower mitochondrial toxicity.

## RESULTS

### Chloramphenicol and linezolid alter the distribution of mitoribosomes along transcripts

We used mitoribosome profiling as an unbiased approach to interrogate the position of mitoribosomes along transcripts with codon-level resolution^15^. In this technique, mitoribosomes from actively-translating cells are isolated, RNase treatment is used to digest the RNA not protected by the ribosomes, and the remaining mitoribosome-protected footprints are sequenced and mapped to the mitochondrial genome. We performed mitoribosome profiling and total RNA sequencing in HEK293 suspension cells subjected to brief (5 minutes) treatment with 300 μM CHL or LZD (or dimethyl sulfoxide (DMSO) as a control). To minimize artifacts from post-treatment effects, we collected the cells using flash filtration and rapid freezing in liquid nitrogen.

Treatment with CHL or LZD led to distinct changes in ribosomal distribution along the length of each of the 11 mitochondrial transcripts (Fig. 1A), although the relative coverage between transcripts was unaltered (Fig. 1B). Relative ribosomal occupancy, defined as codon-level RPM (reads per million), was directly correlated between CHL- and LZD-treated cells (Fig. S1A, Pearson r 0.588 - 0.768), but distinct from DMSO-treated cells (Fig. S1B, Pearson r 0.348 - 0.542). Cells treated with CHL or LZD had more sites with high relative ribosomal occupancy (≥ 1000 RPM) as compared to the DMSO controls (Fig. 1C). Although many of these sites were located near the start of mitochondrial transcripts, the profiles of CHL- and LZD-treated samples were distinct from those of cells treated with the translation initiation inhibitor retapamulin (RET) (Fig. S1B, Pearson r 0.295 - 0.502), indicating that CHL and LZD are inhibiting translation elongation rather than initiation.

**Figure 1:**
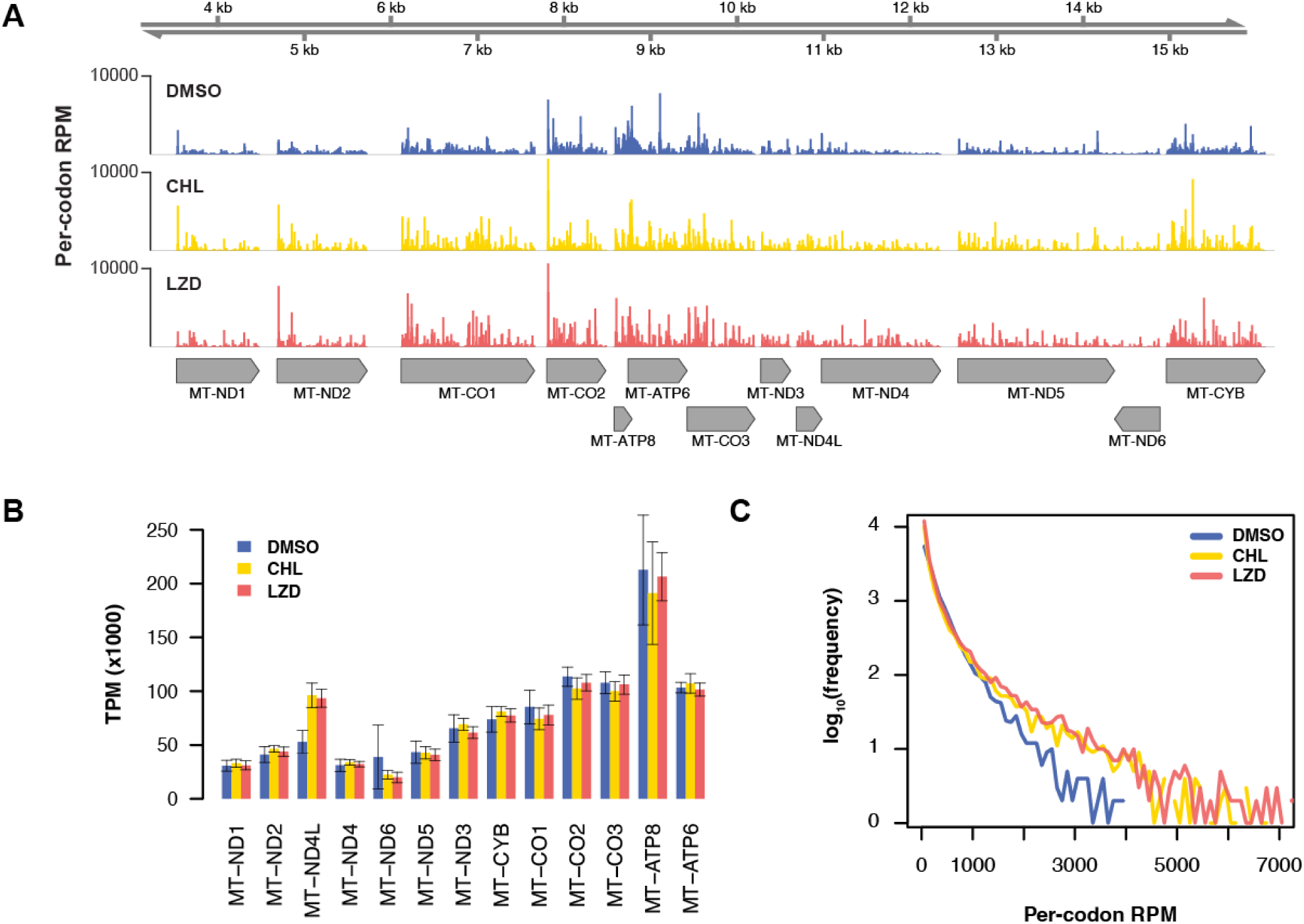
Linezolid and chloramphenicol alter the distribution of ribosomes along each mitochondrial transcript without altering overall coverage. **A)** Per-codon reads per million (RPM) counts for mitoribosome protected fragments (mitoRPFs) for protein-coding genes across the mitochondrial genome. One representative replicate is shown for each sample. **B)** Relative synthesis of mitochondrial gene products across each open reading frame, expressed as transcripts per million (TPM) in cells treated with chloramphenicol (CHL), linezolid (LZD), or dimethyl sulfoxide (DMSO). Error bars indicate standard deviation across replicates. **C)** Mitochondrial transcriptome-wide frequency of per-codon RPM in each treatment condition. Data from all replicates is included.

If CHL and LZD were inhibiting translation elongation generically, the distribution of ribosomes in cells treated with these antibiotics should parallel the distribution of untreated cells. The altered distribution observed therefore argues against such a general mechanism and suggests that inhibition of mitoribosomal translation by CHL and LZD is contextually mediated.

### Chloramphenicol and linezolid show evidence of sequence-specific stalling

To investigate factors mediating antibiotic-induced translational inhibition, we examined ribosome occupancy at the individual codon level. Codon positional occupancy analysis of samples treated with CHL or LZD compared to DMSO controls revealed that the majority of predictive information was contained in the P-1 position (Fig. 2A). Most distinctively, there was a pronounced bias towards alanine in the P-1 position in antibiotic-treated samples. Alanine was present in the P-1 position in roughly a quarter of ribosomal footprints in CHL- and LZD-treated cells (22% and 25%, respectively), as opposed to only 7.1% in DMSO-treated cells. Serine and threonine were also overrepresented, to a lesser extent, in the P-1 position (CHL: 14% S, 12% T; LZD: 11% S, 18% T; DMSO: 7.0% S, 7.6% T) (Fig. 2B). These biases are consistent with those reported from ribosome profiling performed in bacteria^7^. Gly was also enriched in the P-1 position of CHL-treated, but not LZD-treated cells (DMSO: 5.1%, CHL: 8.6%, LZD: 2.9%). The bias towards A/S/T/(G) in the P-1 position was not codon-specific (Fig. 2C), indicating that causative effects are mediated by the amino acid incorporated into the nascent chain, rather than by the corresponding codon or tRNA.

**Figure 2:**
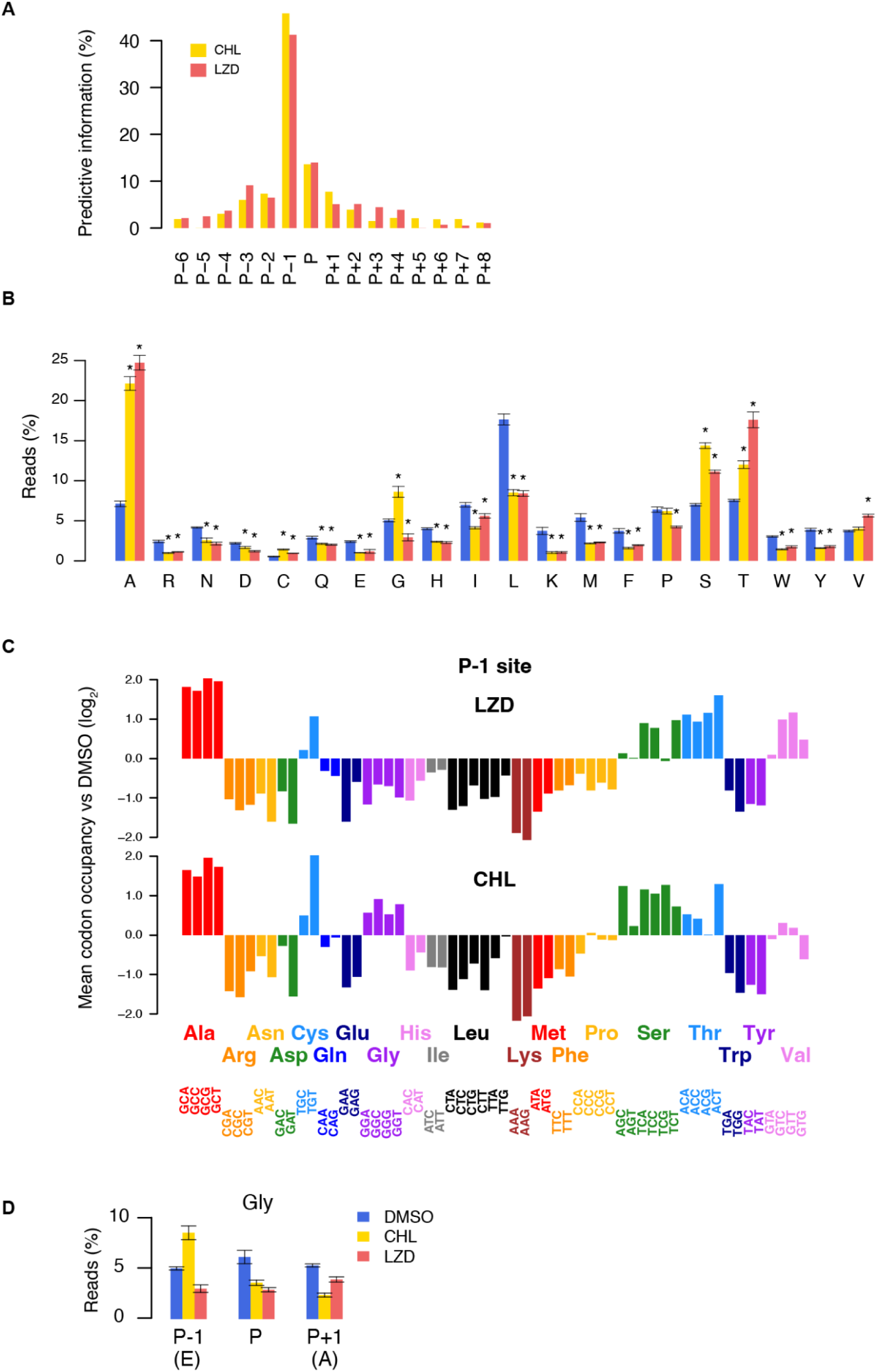
Mitoribosome profiling reveals a strong A/S/T bias in the P-1 position of cells treated with linezolid or chloramphenicol. **A)** Predictive information of each codon position of mitoribosome footprints with respect to the ribosomal P site. Predictive information was calculated from the percentages of reads with each amino acid in each position as in (B) (see Methods). **B)** Percentage of mitoribosome footprint reads with codons for each amino acid in the P-1 position. Error bars indicate standard deviation across replicates. Asterisks indicate a statistically significant difference compared to DMSO (Student’s t-test, p < 0.05). **C)** Mean codon occupancy (see Methods) versus DMSO control at the P-1 site in each treatment condition. Each codon must be present at least four times in the transcriptome to be included. **D)** Percentage of mitoribosome footprint reads with codons for glycine in the P-1, P, and P+1 positions, as in (B). Error bars indicate standard deviation across replicates. LZD: linezolid, CHL: chloramphenicol, DMSO: dimethyl sulfoxide

Although a P-1 alanine was the strongest predictor of stalling, not all instances of alanine led to stalls. Therefore, we further investigated secondary mediators. Our analysis of all footprints revealed an underrepresentation of glycine in the P and P+1 (A) positions in antibiotic-treated cells (Fig. 2D). Gly was present in the P position of footprints 6.2% of the time in DMSO-treated cells, but only 3.6% and 2.9% of footprints in cells treated with CHL or LZD, respectively. Similar decreases in Gly abundance were seen in the P+1 position: 5.4% in DMSO-treated cells, compared to 2.4% in CHL-treated cells, and 3.9% in LZD-treated cells. Pairwise analysis of P-1:P and P-1:P+1 positions revealed that, with CHL, Gly in either the P or the P+1 site appears to alleviate pausing associated with Ala in the P-1 site, although the effect is more pronounced with Gly in the P+1 site (Fig. S2A-B). With LZD, Gly in P site, but not Gly in the P+1 site appears to alleviate such pausing (Fig. S2C-D). Alleviation of stalling by glycine in the P or +1 position is consistent with observations from bacterial systems^7,8^. The lack of steric hindrance of a tRNA-bound nascent peptide containing Gly at its C-terminus and the small size of Gly as an incoming nucleophile have have been proposed to allow for peptidyl transfer even in the presence of nascent peptide-stabilized, PTC-bound antibiotic^8,9^.

To better understand sequence features of nascent peptides that cause the greatest inhibition of mitochondrial translation in the presence of each antibiotic, we next focused our analysis on the most prominent stalling sites. For this purpose, we defined “stall sites” as codons where the ribosomal RPM was > 2-fold over the mean RPM in a 7-codon sliding window (n = 5 datasets for CHL, 6 for LZD and 4 for DMSO). Using this criteria, we identified 321 stall sites in CHL-treated cells, 313 stall sites in LZD-treated cells, and 168 stall sites in DMSO-treated cells (Fig. 3A).

**Figure 3:**
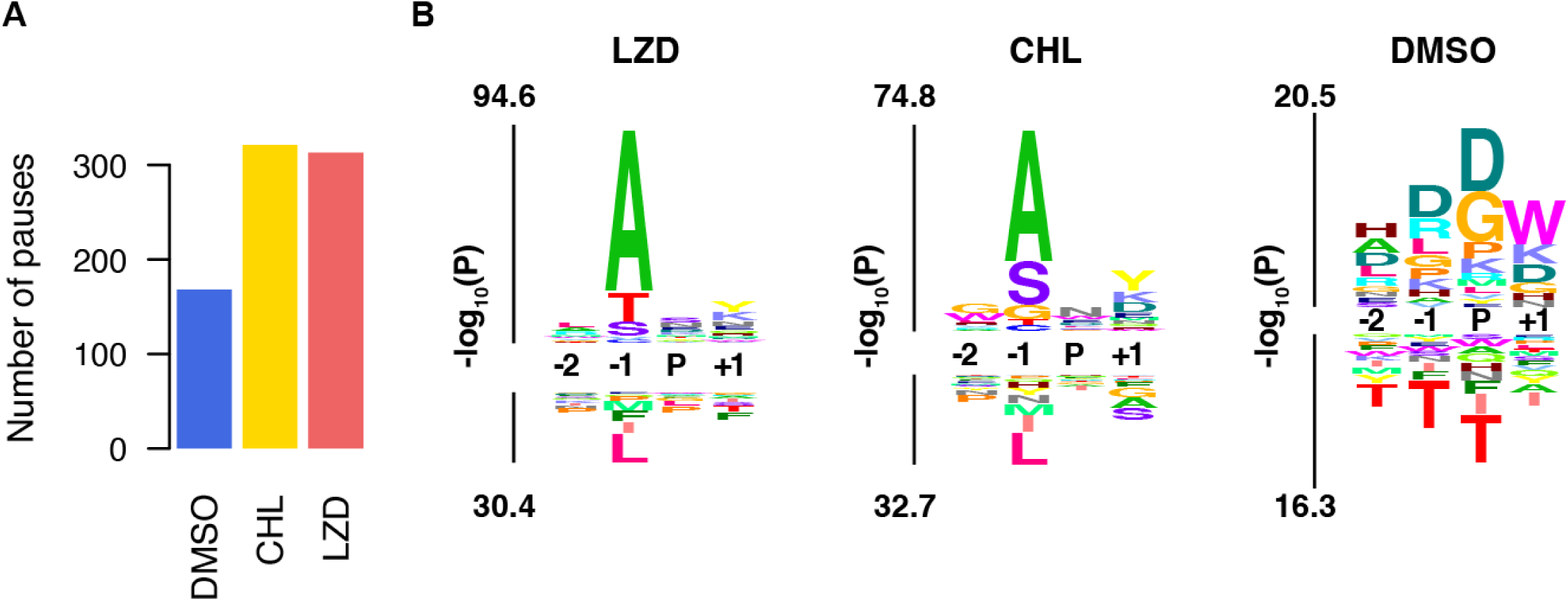
Linezolid and chloramphenicol treatment leads to increased, and biased, ribosomal pausing. **A)** Number of pause sites, defined as codons where the ribosomal reads per million (RPM) is > 2-fold over the mean RPM in a 21-codon sliding window in cells treated with chloramphenicol, linezolid, or DMSO. **B)** kpLogo^3^ probability logo of stall site sequences reveals a large P-1 A/S/T bias in stall sites of LZD- and CHL-treated cells (left and middle), that is absent in those of DMSO-treated cells (right), which are much more heterogeneous. Background was taken as the average across all positions in input sequences. Y axis represents statistical significance determined by one-sided binomial tests of each residue at each position; enriched residues stack on the top, whereas depleted residues stack on the bottom. LZD: linezolid, CHL: chloramphenicol, DMSO: dimethyl sulfoxide

Alanine was the most prominent amino acid present in the P-1 position of these stall sequences (35% of LZD stall sites and 28% of CHL stall sites), followed by threonine (25% in LZD, 15% in CHL) and serine (17% in LZD, 23% in CHL) (Fig. 3B). These values were much higher than those found in DMSO stall sites (8% A, 3% T, 6% S), which were similar to those expected by chance (7% A, 7% S, 9% T). These values must be interpreted cautiously, however, due to the limited numbers of potential sequences. When we reanalyzed published mitoribosomal profiling datasets from HeLa cells treated with CHL^15^, we found a highly similar stalling profile (Fig. S3).

### In vitro reconstitution validates critical role of −1 amino acid in translation inhibition

To evaluate the ability of CHL and LZD to directly cause sequence-specific stalling of mitoribosomes, we performed in vitro translation in a fully reconstituted mitochondrial translation system^16^. This system allows us to probe the dependence of stalling on specific amino acid residues within the nascent chain and test the ability of our mitoribosome profiling to accurately identify mitoribosome stalling directly caused by antibiotics. To this end, we designed a split nanoluciferase (nLuc) assay with a reporter transcript consisting of a putative stall sequence followed by an 11-amino acid HiBiT peptide^17,18^(Fig. 4A). In the presence of the complementary LgBiT protein, active nLuc is formed, leading to the production of luminescence when supplied with an appropriate substrate. The short length of the HiBiT, as well as the lack of alanines in the sequence, minimizes the chance of antibiotic-induced stalling during its translation (Supplementary Table 1). This allows luminescence to serve as a read-out of translation of the preceding test sequence. Furthermore, in order to minimize potential tRNA-dependent artifacts, we used human in vitro transcribed tRNAs for incorporation of the P and P+1 (A-site) amino acids in the test sequence (Supplementary Table 2).

**Figure 4:**
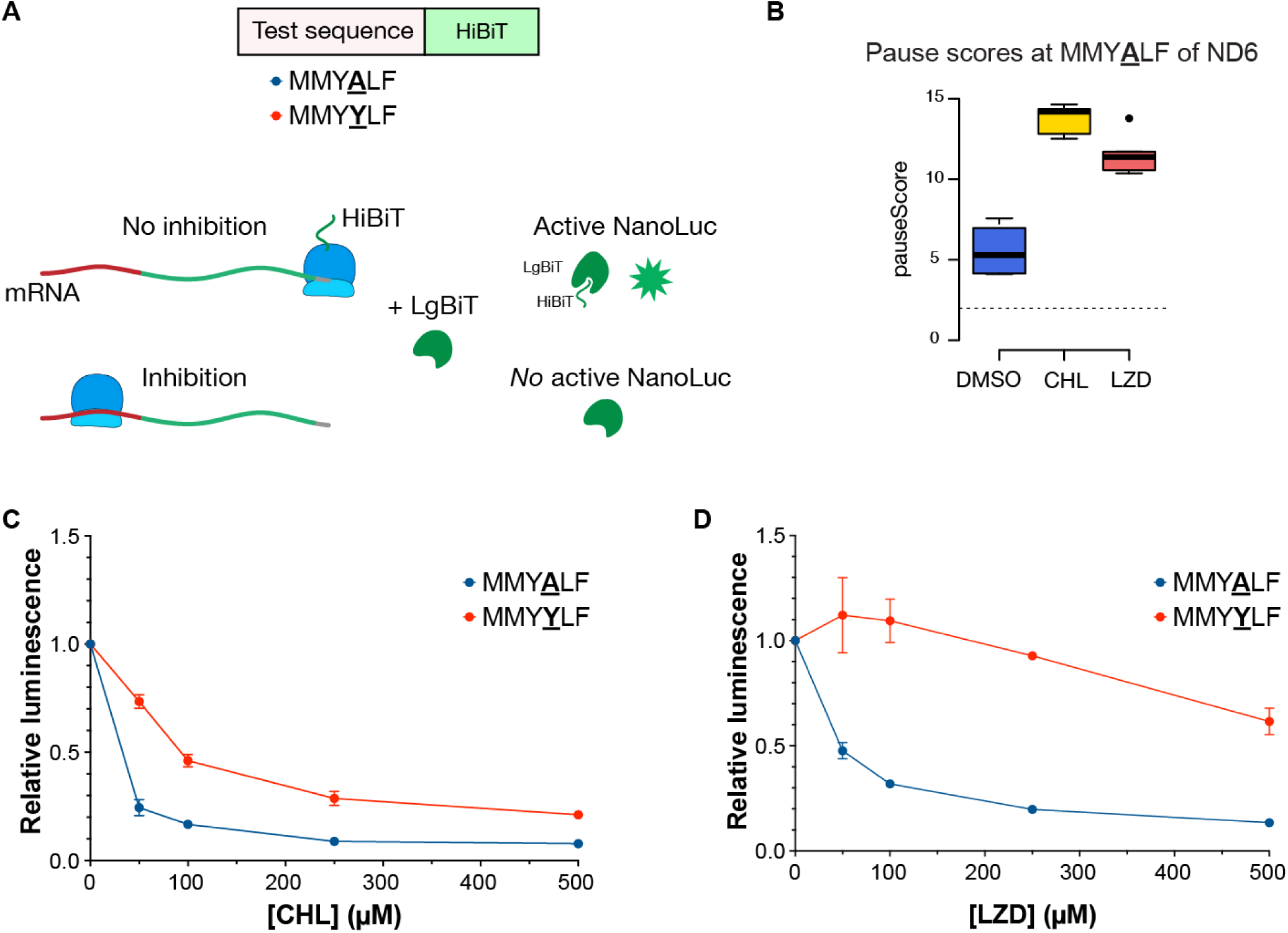
A stall site identified by mitoribosome profiling produces sequence-specific stalling by chloramphenicol and linezolid in a mitochondrial in vitro translation system. **A)** Schematic of HiBiT-based reporter assay. **B)** Box-and-whisker plot of pause scores (fold-changes over the mean of a 21-codon window, see Methods) for the ND6 sequence MMY**<u>A</u>**LF (P-1 site underlined). The dotted line at two shows the cutoff for pause-calling. **C-D)** Both linezolid (LZD) **(C)** and chloramphenicol (CHL) **(D)** robustly inhibited translation of the identified stall sequence MMYALF in a dose-dependent manner (blue). Significantly weaker inhibition was seen when the P-1 position (underlined and bolded in **A**) was changed from an alanine to a tyrosine (red), consistent with the A/S/T bias observed through mitoribosome profiling. Luminescence values are plotted relative to dimethyl sulfoxide (DMSO) control. Error bars indicate standard error of the mean (SEM). n = 3 technical replicates.

For a test sequence, we chose a mitochondrial stall site near the beginning of ND-6, MMY<u>A</u>LF (P-1 site underlined). This site had median pause scores (fold-change over the mean of a 21-codon window, see Methods) of 11 in LZD-treated cells and 14 in CHL-treated cells, as compared to only 5 in DMSO-treated cells (Fig. 4B). It also contained only a single A/S/T residue, which would allow for unambiguous interpretation of results, and had a sequence that was compatible with current limitations in the in vitro aminoacylation of human mitochondrial tRNAs. Both CHL and LZD inhibited translation of the MMYALF reporter in a concentration-dependent manner, with IC50 values of 13 μM (95% confidence interval 9.5-17 μM) for CHL (Fig. 4C) and 38 μM (30-47 μM) for LZD (Fig. 4D). This inhibition was strongly dependent on the identity of the amino acid in the P-1 position; changing the P-1 alanine to a tyrosine led to a nearly 8-fold increase in the IC50 of CHL (99 μM (70-125)), and increased the IC50 of LZD to > 500 μM (Fig. 4C, D).

These results further support that CHL and LZD do not act as general elongation inhibitors; rather, the ability of these antibiotics to inhibit translation is dependent on the amino acids in the sequence that is being translated. Furthermore, these data validate the ability of our mitoribosomal profiling data analysis to identify sequences conducive to CHL- and LZD-induced stalling. The luminescent readout only informs on extent of inhibition and not location of stalled ribosomes; however, in combination with the profiling data, these in vitro translation assay results are consistent with a model in which an alanine in the P-1 position facilitates CHL- and LZD-induced mitoribosomal stalling.

### The structure of the human mitoribosome bound to linezolid reveals a highly conserved binding mechanism

The above model is consistent with observations made in bacterial ribosome inhibition, where the nascent peptide contributes to the formation of an antibiotic binding site within the ribosome^7–10^. The nascent peptide-dependent antibiotic binding in bacteria is a consequence of steric accommodation, as well as stabilizing interactions between the antibiotic and the amino acid residue in the P-1 position. More specifically, the sequence of the nascent chain shapes the antibiotic binding site, with the P-1 amino acid residue either occluding (in the case of large amino acids) or forming stabilizing interactions (in the case of A/S/T) with the antibiotic. In contrast, little is known about binding mode of these antibiotics to mitochondrial ribosomes, apart from cross-linking of a photoactivatable LZD derivative implicating a similar binding site in the PTC^6^.

We purified 55S mitoribosomes (mitomonosomes) from FreeStyle^TM^ 293-F cells (HEK293 cells adapted for serum-free suspension culture) subjected to a brief, high concentration treatment with linezolid (5 min at 100 μg/μL (300 μM)) (see Methods). We verified the assembly of the 55S mitoribosome through 2D classification (Fig. S4). Upon 3D reconstruction, we observed significant disordering and/or dissociation of the small (28S) subunit (mt-SSU) relative to the large (39S) subunit (mt-LSU). Since crosslinking data^6^ and homology to the bacterial ribosome suggest that LZD binds in the PTC in the mitochondrial LSU, we proceeded with data collection and processing, determining a 2.62 Å cryo-EM structure of the large subunit of the human mitoribosome bound to linezolid (Figs. S5 and S6, Supplementary Table 3). The density in the PTC is particularly well resolved, permitting the modeling of the conformation of the drug and surrounding solvent molecules.

To determine how the mitoribosome binds to LZD, we compared our LZD-bound structure to a previously determined apo mitoribosome (PDB 7QI5^19^) (Fig. 5B). As expected, the conformations of most of the nucleotides in the PTC were unperturbed by LZD. A notable exception is U2993 of the 16S rRNA, whose ribose is shifted away to accommodate the bulky fluorophenyl B ring of LZD, avoiding a clash (Fig. 5B). We also observe density consistent with a metal, likely Mg^2+^, that is coordinated, in part, by the oxazolidinone moiety of the antibiotic (Fig. 5B). Both of these features are consistent with the features observed in our previous structures of the *E. coli* ribosome bound to LZD alone (PDB 7S1H) and LZD with a stalled peptide (PDB 7S1G): the analogous base (U2506 in *E. coli*) shifts to accommodate LZD and a metal binds the oxazolidinone ring with similar geometry^10^.

**Figure 5:**
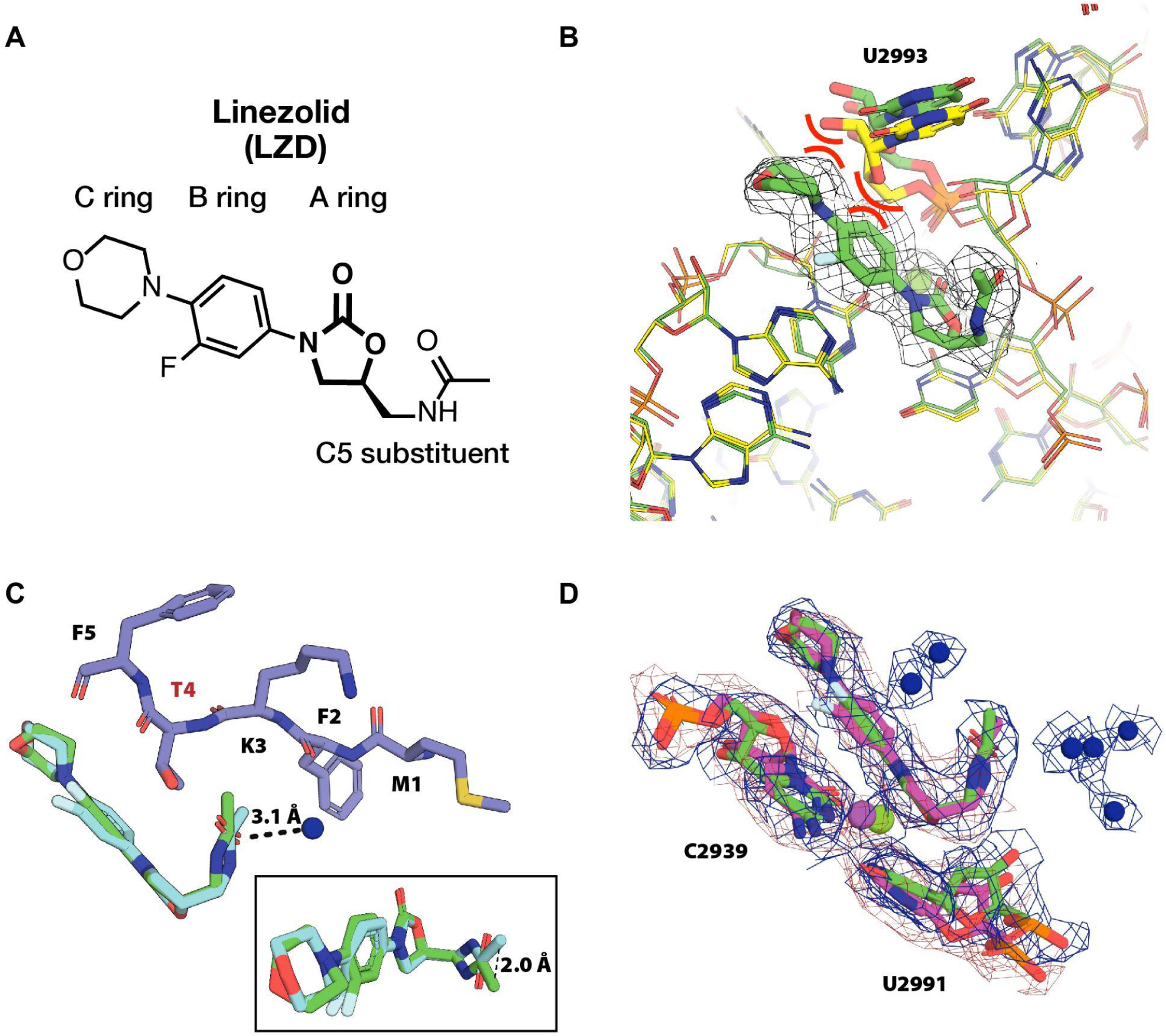
Structure of the large (39S) subunit of the human mitochondrial ribosome (mt-LSU) in complex with linezolid shows strong water density in the peptidyl transferase center (PTC) and an ordered C5 side chain. **A)** Chemical structure of linezolid (LZD), with key structural features indicated. The oxazolidinone (A) ring, characteristic of oxazolidinone antibiotics, is shown in bold. **B)** Structure of LZD in the mt-LSU (present paper, green) compared to the apo mitoribosome (PDB 7QI5^1^, yellow). LZD clashes with the ribose of U2993 of the 16S rRNA in the apo structure, necessitating a conformational adjustment. Metal ion, assigned as Mg^2+^, interacts with the carbonyl oxygen of the A-ring and ribosomal RNA. **C)** Overlay of the mitoribosome-bound LZD (this paper, green) and the stalled complex of LZD and a nascent chain on the *E. coli* ribosome (PDB 7S1G^2^: bacterial ribosome nucleotides, cyan and nascent chain, purple). The C5 tail conformation is altered by displacement and a torsional change that displays the amide carbonyl towards the nascent chain. A threonine residue has been modeled in place of the alanine present in the nascent chain observed in the structure in order to demonstrate the possible accommodation of the bulkier residue in the mitoribosomal structure given the altered C5 conformation. A hydrogen bond of the carbonyl group of the amide to a solvent molecule (blue sphere) is indicated by dashed lines. **D)** Density for solvent molecules in the LZD-bound PTC of the mitoribosome observed at 3.8 **σ**. No solvent molecule density is observed in the LZD-bound structure of the *E. coli* ribosome (PDB 7S1H, magenta) at 2.2 **σ**, which equivalently encompasses the surrounding bases, LZD, and a metal ion, assigned as Mg^2+^.

To directly compare the bacterial and mitochondrial binding modes, we aligned the mitoribosome-LZD complex to the structure of the *E. coli* LZD-stalled ribosome complex (PDB 7S1G) using the LZD A, B, and C ring atoms (RMSD 0.48 Å) (Fig. 5C). In *E. coli* ribosomes, the density supporting the conformation of the C5-tail of LZD was stronger for the stalled complex than the otherwise empty ribosome; nevertheless, we observe strong density supporting modeling of the C5-tail conformation of the mitoribosome complex even in the absence of a nascent chain. In the mitoribosome, the conformation of the LZD C5-tail moves 2.2 Å away from the position observed in the stalled *E. coli* complex. In addition, a torsional change repositions the amide carbonyl group of the C5 tail to face where the putative nascent chain would be located. Although, as expected due to dissociation of the small subunit, we do not observe any density for a nascent chain in the mitoribosome complex, we do observe strong density for solvent molecules hydrogen bonded to the amide moiety of the C5 tail. In addition, the twisted conformation of the C5 tail alters the “binding pocket” for the nascent peptide. Its pivot away may explain the relaxed specificity for accommodating the bulkier Thr residue in the P-1 position for the mitochondrial ribosome compared to *E. coli* (Fig. 5C). This conformational preference may be driven by the changes in solvation patterns observed between the two ribosomes. Despite worse global resolution for our mitoribosome-LZD complex (2.62 Å) compared to our previous LZD-*E. coli* ribosome complex (2.35 Å), we observe stronger density for solvent molecules throughout the PTC of the mitoribosome (Fig. 5D, S7). This suggests that the ability of the C5 tail of LZD to interact with nascent chains may differ when bound in the two PTCs, and indicates that considering solvent interactions could provide an important selectivity handle in the design of oxazolidinone derivatives with reduced mitoribosome toxicity.

## DISCUSSION

The use of chloramphenicol and oxazolidinone antibiotics as antibacterial therapies has been limited by host mitotoxicity caused by off-target inhibition of mitochondrial translation^4,20^. For example, linezolid (LZD) treatment is frequently discontinued due to reversible hematologic abnormalities^21–23^. Here, through mitoribosome profiling and in vitro translation assays, we show that these antibiotics cause context-specific stalling of mitoribosomes along transcripts in a manner consistent with their effects on bacterial ribosomes. Together with our cryo-EM analysis of the human mitoribosome bound to LZD, these results provide critical insights into mode of inhibition of the mitoribosome by LZD, and may help inform the development of rationally-designed next-generation oxazolidinone antibiotics. Such antibiotics are in increasingly high demand as global levels of antimicrobial resistance rise^1^.

Our mitoribosome profiling data revealed that mitoribosomes become strongly, and reproducibly, stalled at discrete locations along each transcript in the presence of CHL and LZD (Fig. 1). Examination of these locations revealed a strong bias towards the presence of an alanine (and to a lesser extent serine or threonine) in the −1 position with respect to the P site, as well as a bias against glycine in the P and P+1 (A) sites (Fig. 2)^6^. We were able to recapitulate antibiotic-dependent stalling at one such identified site in a fully reconstituted in vitro mitochondrial translation system (Fig. 4). We further confirmed the importance of the P-1 amino acid for this stalling, as substitution of a P-1 alanine to a tyrosine dramatically decreased the ability of LZD and, to a lesser but still significant extent, CHL, to inhibit translation of this sequence. In bacteria, this is due to mutually exclusive binding of antibiotic and a large amino acid at the P-1 site^7–9^. Our findings indicate a mechanism of inhibition for CHL/LZD-mediated mitoribosome stalling consistent with that observed in bacteria, whereby the identity of the P-1 amino acid determines whether the antibiotic can stably bind to the peptidyl transfer center.

Our reanalysis of published mitoribosomal profiling datasets from human cells treated with CHL identified a highly similar stalling profile (Fig. S3). This is in contrast to a recent manuscript reporting a lack of sequence bias caused by CHL in mitoribosome profiling experiments^24^. In said study, however, CHL was only included in the lysis buffer. This is in contrast to our experimental design, in which cells were pretreated with CHL prior to harvesting. Given the inefficiency of mitochondrial translation and the extreme dilution upon lysis, lack of pre-treatment could account for the apparent discrepancy. As our study is the first report of mitoribosome profiling performed with LZD, no previous codon-level mitoribosome translation inhibition data is available for this antibiotic.

In bacterial ribosome profiling, a stronger alanine bias was found in LZD-compared to CHL-treated cells, with serine and threonine showing increased representation in the P-1 position in CHL-treated as opposed to LZD-treated cells^7^. This was attributed to the presence of an additional nascent peptide-antibiotic interaction between the hydroxyl group of threonine (and presumably serine) and one of CHL’s chlorines^9^. Our mitoribosome profiling similarly showed a stronger alanine bias in the P-1 position in LZD-, compared to CHL-treated cells (Fig. 2B); however, we saw relatively increased representation of serine and threonine in the LZD-treated cells compared to the bacterial findings. This Thr enrichment in LZD-treated cells suggests that there may be subtle differences in the binding interactions between the nascent chain and LZD in mitochondrial ribosomes compared to bacterial ribosomes.

To investigate the structural basis of context-dependent stalling of mitoribosomes, we obtained the cryo-EM structure of the large subunit (LSU) of the human mitoribosome bound to LZD (Fig. 5). Comparison with LZD bound to a bacterial ribosome revealed that the binding of LZD to mitoribosomes is similar to its binding to bacterial ribosomes, with notable differences in the conformation of the C5 tail and the extent of solvation in the PTC when LZD is bound (Fig. 5). The altered conformation of the tail may underlie the higher amount of stalling on Thr observed for the mitochondrial ribosome relative to the *E. coli* ribosome. Resolving a stalled complex of a Thr-containing nascent peptide chain stalled on the mitoribosome will be needed to dissect the structural basis of this change in specificity. The unique features of mitochondrial translation and the extensive process of mitoribosome purification, however, present steep challenges to its ascertainment.

Our present structure can inform strategies for widening the selectivity window between bacterial and mitoribosomes, which is one of the barriers limiting the wider clinical use of oxazolidinone antibiotics. Increased solvation of the PTC region of the mitoribosome, as compared to the *E. coli* ribosome, influences the conformation of the C5 tail of LZD. Importantly, the ordering of water goes against expectation based on the global resolution of the density maps: the higher-resolution *E. coli* map has fewer ordered waters. Collectively, these results point to the importance of considering solvation in tuning selectivity for highly conserved binding sites^25,26^.

In summary, our findings indicate that inhibition of mitoribosome translation by CHL and LZD antibiotics occurs in a nascent peptide sequence-selective manner, a feature conserved with inhibition of the bacterial ribosome by these compounds. Our study lays the groundwork for future, targeted, investigation into selectivity and mitotoxicity of oxazolidinone compounds and provides a high-resolution structure of the binding site of linezolid on the human mitoribosome that can serve as a valuable resource in the design of next-generation antibiotics.

### Limitations of this study

In this work, we carried out mitoribosome profiling in a single cell type, FreeStyle^TM^ 293-F. We uncovered evidence of similar CHL–associated mitotranslational biases in an independent dataset collected in HeLa cells^15^ (Fig. S3); however, future studies in additional cell types will be needed in order to determine the broader generalizability of our findings, especially with regards to LZD, for which no other mitoribosome profiling data is available for comparison. Nevertheless, the ability to recapitulate an identified stall site in a fully reconstituted system indicates that the core translational machinery is sufficient to mediate the observed context-specificity. Due to current technical limitations of the in vitro translation system, we were unable to further interrogate the determinants of mitoribosomal stalling, including the ability of a glycine in the P and/or P+1 positions to overcome stalling. Human mt-tRNAs have distinctive structures^27,28^ and many, including that for glycine, pose currently insurmountable challenges to in vitro transcription and aminoacylation. This highly restricts the choice of test sequences for the HIBiT assay. Advances in the reconstituted transcription and translation systems, namely the ability to generate and aminoacylate all human mt-tRNAs, could allow for testing additional sequences to further investigate our proposed model.

Although the concentrations of antibiotics we used for footprinting (300 μM (~100 µg/mL)) were in line with those commonly used in the laboratory, they were well above levels cells are likely exposed to during the course of antibiotic treatment – serum LZD levels in one study were found to average 13.4 μg/mL^29^. Our treatment, however, was also much briefer than clinical courses of LZD treatment, which can last 6 months in the case of some drug-resistant tuberculosis^30^. It is unknown how our results carry over to a lower-dose, longer-period, treatment time.

Additionally, we did not investigate any physiological effects of sequence-selective inhibition. The stall sites we observed through mitoribosome profiling were distributed over all transcripts (Fig. 1A). However, it is possible that, similarly to the distinct phenotypic effects of different mtDNA mutations, preferential stalling of specific transcripts could lead to distinctive respiratory chain imbalances.

## Acknowledgements

We thank Stephen Floor, Albert Xu, and Yan Zhang for guidance with ribosome profiling; Giovanni Aviles for assistance with mitoribosome purification and mitoribosome profiling; and Daniel Hogan for help with data storage and transfers. We also thank Xiaoyang Guo and Mo Yao for assistance with in vitro translation assays and Annía Rodriguez Hernández, Chloe Ghent, and Ada Álvarez Muñoz for helpful discussions. We acknowledge support from NIAID (R01AI137270 to D.G.F.), NIGMS (R01GM123002 to L.S.C.), the W.M. Keck Foundation Medical Research Grant (to J.S.F. and D.G.F.), the UCSF Discovery Fellowship (to J.I.K.), and Grants-in-Aid for Scientific Research from JSPS (23H04252 to N.T.).

## Author contributions

B.B. conceived the research, designed and performed mitoribosome profiling and purification of mitoribosomes for structural characterization, analyzed data, and wrote the manuscript. T.R. performed structural analysis and assisted in manuscript writing. M.C. analyzed mitoribosome profiling data and assisted in manuscript writing. MH performed in vitro translation assays and assisted in data interpretation. J.I.K assisted in experimental design, data interpretation, and manuscript editing. N.T. supervised the in vitro translation experiments, assisted in experimental design and data interpretation, and edited the manuscript. L.S.C. supervised the mitoribosome profiling data analysis, assisted in data interpretation, and edited the manuscript. J.S.F. supervised the structural analysis, assisted in experimental design and data interpretation, and assisted in manuscript writing. D.G.F. conceived and supervised the research, assisted in data interpretation, and edited the manuscript.

## Data availability

Atomic coordinates for the structure of the human mitochondrial ribosome large subunit (39S) bound to linezolid has been deposited in the PDB under the accession number 9CN3 and the EMDB under the accession number EMD-45757. Raw and processed sequencing data were deposited in the GEO database under the accession number GSE273206. Processing and analysis scripts are available at GitHub (https://github.com/churchmanlab/human-mitoribosome-profiling_appendixA : an add-on to https://github.com/churchmanlab/human-mitoribosome-profiling).

## METHODS

### Mitoribosome profiling

#### Cell culture and treatment

FreeStyle^TM^ 293-F cells (HEK293 cells adapted for serum-free suspension culture) were grown in suspension culture at 37°C, 5% CO_2_, with orbital shaking (115 rpm) in unmodified FreeStyle^TM^ 293 Expression Medium (Gibco) in the absence of antibiotics. Cell cultures were diluted to 2 x 10^6^ viable cells/mL and 50 mL portions were transferred into 125 mL flasks before use. To each portion, DMSO or antibiotic was added to a final concentration of 1% DMSO, 300 μM (97 μg/mL) chloramphenicol (CHL), 300 μM (101 μg/mL) linezolid (LZD), or 100 μM (155 μg/mL) retapamulin (RET). Samples were placed back into the incubator for 5 minutes before collecting cells via fast vacuum filtration through a 0.8 μM filter. The collected cells were then quickly scraped off the filter and plunged into liquid nitrogen. The frozen cells were stored at −80°C until use.

#### Mitoribosome footprint isolation

Mitoribosome footprints were prepared as described^15^ with modifications. Each 10 x 10^6^ cell pellet of frozen cells was resuspended in 900 μL of mito-RP lysis buffer (10mM Tris pH 7.5; 50mM NH_4_Cl, 20mM MgCl_2_, 0.25% lauryl maltoside, 10mM DTT, 1X EDTA-free protease inhibitor cocktail (Pierce) containing antibiotic or DMSO at the same concentration as in the pre-treatment) and incubated for 10 min on ice. It was then homogenized via 15 strokes of a Dounce homogenizer and centrifuged for 1 min @ 1000 x g. The supernatant was transferred to new tubes and a 25 μL portion of each was flash frozen in liquid nitrogen for total RNA sequencing. RNase If (NEB) was added to each supernatant for a final concentration of 7.25 U/μL to produce mitoribosome footprints. Samples were incubated for 30 minutes at room temperature without rotation. 80 U of SUPERaseIn^TM^ RNase Inhibitor (Invitrogen) was then added per 450 μL of digested lysate to prevent overdigestion and the digested lysate was clarified by centrifugation in a benchtop microcentrifuge @ 10,000 RPM, 5 min, 4°C. Clarified lysates were then transferred to a 10-50% linear sucrose gradient (prepared in mito-RP lysis buffer without lauryl maltoside & protease inhibitor, using a BioComp GradientMaster^TM^) and centrifuged for 3 h @ 4°C, SW41Ti rotor. Fractions were collected with a BioComp Piston Gradient Fractionator^TM^ and portions corresponding to mitomonosomes (identified based on UV 260 trace and confirmed via western blot for MRPL12 and MRPS18B) were frozen @ −80°C in 3 volumes of TRIzol^TM^ LS (Invitrogen). Footprints were then isolated using a Direct-zol^TM^ RNA kit (Zymo) and subsequently concentrated using a RNA Clean and Concentrator-5 kit (Zymo). RNA was then size-selected by running a 15% TBE-Urea gel and extracting fragments between 28 and 40 nt in length, corresponding to mitoribosome-protected footprints. DMSO: n = 4 biological replicates (separate cell cultures), CHL: n = 4 biological replicates and 1 technical replicate (separate datasets from same culture); LZD: n = 4 biological replicates and 2 technical replicates; RET: n = 2 biological replicates.

#### Sequencing library preparation

Libraries were prepared from the extracted footprint RNA as described^31–34^ with modifications. Briefly, extracted RNA was desalted using a RNA Clean and Concentrator-5 kit (Zymo), then dephosphorylated with T4 PNK (NEB) and ligated to a preadenylated oligonucleotide linker (NI-816: 5’-/5Phos/NNNNNTAGACAGATCGGAAGAGCACACGTCTGAA/3ddC/-3’) using T4 RNA Ligase 2 truncated KQ (NEB). Unligated linker was depleted by treatment with yeast 5’-deadenylase (NEB) and RecJ exonuclease (Lucigen/Epicenter), and the linker-ligated footprints were then purified via a Oligo Clean and Concentrator kit (Zymo). The ligation products were then reverse transcribed using Protoscript II reverse transcriptase (NEB) and NI-802 primer (5’-/5Phos/ NNAGATCGGAAGAGCGTCGTGTAGGGAAAGAG/iSp18/GTGACTGGAGTTCAGACGTGTGCTC-3’). The RNA template was then hydrolyzed with NaOH. The reverse transcription products were desalted and concentrated with an Oligo Clean & Concentrator kit (Zymo), followed by gel purification via a 15% TBE-Urea PAGE gel. Extracted cDNA was then circularized using CircLigase II (Lucigen). Circular DNA was quantified via qPCR, then amplified via PCR with the forward primer NI-798 5′-AATGATACGGCGACCACCGAGATCTACACTCTTTCCCTACACGACGCTC-3’ and a unique reverse indexing primer (containing a unique 6 nucleotide barcode) for each sample. PCR products were then purified via a DNA Clean & Concentrator 5 column (Zymo) followed by gel purification via an 8% non-denaturing TBE PAGE gel. Extracted DNA was then quantified and checked for quality with a BioAnalyzer (Agilent).

#### RNA sequencing

Equimolar concentrations of all samples were pooled and sequencing was performed by the UCSF Center for Advanced Technology on an Illumina HiSeq 4000 (SE65 66x8x8x0).

#### Mitoribosome profiling data analysis

Mitoribosome profiling data was processed as described in^35^. Reads in the size range 33 to 39 nucleotides were used for A-site transformation and downstream steps. Coverage across each codon was calculated by summing the read counts across the three sub codon positions and counts were normalized to reads per million mitochondrial mRNA-mapped reads (RPM). Relative synthesis of mitochondrial gene products was determined using Rsubread featureCounts^36^ as described^4^. All read sizes were used and values were normalized by ORF length and total transcript counts (transcripts per million, TPM).

Predictive information at each ribosomal position (e.g. P-1, P, P+1 etc.) was calculated from the percentage of reads with each amino acid in each position. First, data from plots like that in Fig. 1B was collected for all ribosomal positions P-6 to P+8. For each, the absolute difference between the percentage of reads in the treatment condition and in DMSO was taken for all amino acids and summed. Next, the minimal value (P-5 for CHL and P+5 for LZD) was taken as background and subtracted from each other value. Finally, predictive information across these 15 positions was set to 100% and each expressed as a percentage.

Relative ribosome occupancies for codons (as in Fig 2C) were computed by taking the ratio of the A-site transformed ribosome density in a 3-nt window at the codon over the overall density in the respective coding sequence. For each codon, the values of all occurrences across the genome were averaged.

Pause scores were calculated as the fold-change over the mean of rpm-normalized values in a 21-codon sliding window. A pause was called on a codon if the pause score was greater than two in all replicates. Pausing peptide motifs were created with the kpLogo^37^ web app (http://kplogo.wi.mit.edu/), used with default parameters (unweighted, background probability calculated by averaging across all positions). Probability logos are shown, in which residues are scaled relative to the statistical significance determined by one-sided binomial tests (−log10(*P* value)) of each residue at each position. Enriched residues stack on the top, whereas depleted residues stack on the bottom.

Processing and analysis scripts are available at GitHub (https://github.com/churchmanlab/human-mitoribosome-profiling_appendixA: an add-on to https://github.com/churchmanlab/human-mitoribosome-profiling).

### Reconstituted mitochondrial in vitro translation assay

#### Preparation of aminoacylated-tRNA mixture

The aminoacylated iVT tRNA_mix_ included 12 iVT tRNA species (Table 2). To prepare the aminoacylated iVT tRNA_mix_, each iVT tRNA species (0.0019 A_260_ units) was first aminoacylated using the cognate charging enzyme and amino acid individually. Thereafter, all aminoacylated iVT tRNAs were combined and subjected to one NAP-5 column, as described^38^. IVT yeast tRNAs, except tRNA^Lys^, were aminoacylated using yeast S100 extracts. Yeast iVT tRNA^Lys^ was aminoacylated utilizing the recombinant *E. coli* lysyl-tRNA synthetase. IVT mt-tRNA^Met^, mt-tRNA^Leu^, and mt-tRNA^Ser^ were aminoacylated by the cognate recombinant mitochondrial aminoacyl-tRNA synthetase, respectively. IVT mt-tRNA^Phe^ was aminoacylated utilizing the recombinant *E. coli* Phenylalanyl-tRNA synthetase.

#### In vitro translation

In vitro translation using a reconstituted mammalian mitochondrial translation system was performed as described^39^. Briefly, the translation mixtures (5 µL) contained 50 mM Hepes-KOH [pH 7.5], 100 mM potassium glutamate, 11 mM Mg(OAc)_2_, 0.1 mM spermine, 1 mM DTT, 0.15 mM of each amino acid (except methionine and cysteine), 0.05 mM methionine, 0.1 mM cysteine, 1 mM ATP, 1 mM GTP, 20 mM creatine phosphate, 0.1 µg 10-formyl-5, 6, 7, 8-tetrahydrofolic acid, 100 nM creatine kinase, 20 nM myokinase, 15 nM nucleoside-diphosphate kinase, 15 nM pyrophosphatase, 0.5 µM IF-2mt, 1.0 µM IF-3mt, 5 µM EF-Tumt, 1 µM EF-Tsmt, 0.5 µM EF-G1mt, 0.5 µM EF-G2mt, 0.5 µM RF-1Lmt, 0.5 µM RRFmt, 5 μM methionyl-tRNA transformylase (*E. coli* MTF), 0.2 µM 55S ribosome, aminoacyl-tRNA mixture (0.045 A_260_ units iVT tRNA_mix_), 0.2 μM mRNA (Table 1), 0.06 μg/μL LgBiT, and indicated concentration of antibiotics. The reaction mixture was incubated at 37°C for 30 min. The nLuc activities were analyzed using 2 µL of sample as described^39^. Under these conditions, the nLuc activities were detected in a linear range over time, and depended on the concentration of the mRNA. IC50 values were determined using nonlinear regression in GraphPad Prism.

### Mitoribosome structural analysis

Human 55S mitomonosomes were purified from linezolid-treated FreeStyle^TM^ 293-F cells using a modification of ^40^ and ^19^, as described below.

#### Cell culture and harvesting

FreeStyle^TM^ 293-F cells (Gibco) were grown in suspension culture at 37°C, 5% CO_2_, with orbital shaking (115 rpm), in unmodified FreeStyle^TM^ 293 Expression Medium (Gibco) in the absence of antibiotics. In total, mitomonosomes from 4L of cells were purified and combined (from 2, 2L purifications) for cryoEM sample preparation. For each 2L preparation, cultures were scaled up to 2L (in 4 x 500 mL portions in 2L flasks) and grown to a final density of ~3 x 10^6^ cells/mL. At that time, cells were treated with a brief (5 min) high concentration of linezolid (100 μg/mL)(300 μM). Cultures were then harvested by centrifugation at 1,000 × g for 7 min, 4°C. The supernatant was decanted and the pelleted cells were washed to remove excess media by resuspension in ice-cold PBS containing a lower concentration of linezolid (50 μg/mL)(150 μM) and EDTA-free protease inhibitor (Pierce). The resuspended cells were centrifuged at 1,200 × g for 10 min, 4°C, after which the supernatant was decanted. The pellets were then weighed and flash frozen in liquid nitrogen before storing at −80°C until use.

#### Mitochondrial purification

Mitochondria from 2 L worth of FreeStyle^TM^ 293-F cells were purified at a time. LZD was maintained in all purification buffers at a concentration of 50 μg/mL (150 μM). Pellets were thawed on ice and resuspended in 60 mL ice-cold MIB buffer (50 mM HEPES-KOH, pH 7.4; 10 mM KCl; 1.5 mM MgCl_2_; 1 mM EDTA; 1 mM EGTA; 1 mM DTT; EDTA-free protease inhibitor (Pierce); 50 μg/mL LZD). The resuspended cells were then incubated at 4° for 15 min with gentle agitation on a nutator. 20 mL SM4 (280 mM sucrose; 840 mM mannitol: 50 mM HEPES-KOH, pH 7.5; 10 mM KCl; 1.5 mM MgCl_2_; 1 mM EDTA; 1 mM EGTA; 1 mM DTT; EDTA-free protease inhibitor (Pierce); 50 μg/mL LZD), equivalent to ⅓ of the MIB buffer volume, was then added and the sample was homogenized via 60 strokes of a Dounce homogenizer (processed in two portions in a 100 mL Dounce). Differential centrifugation was then performed to fractionate the subcellular organelles as described^40^, but with the inclusion of 50 μg/mL LZD (150 μM) in all buffers. Briefly, the homogenized sample was centrifuged at 800 x g for 15 min, 4°C, and the supernatant was filtered through Miracloth. The remaining pellet was resuspended in 20 mL MIBSM buffer (consisting of a 3:1 ratio of MIB:SM4), homogenized by 15 strokes of the Dounce, and centrifuged for 15 min @ 800 x g, 4°C. The resultant supernatant was passed through Miracloth and combined with the previous supernatant. The combined supernatants were then centrifuged at 1,000 x g for 15 min, 4°C. The supernatant from this was then centrifuged for 15 min at 10,000 x g, 4°C. The resultant supernatant and loose pellet were carefully removed, keeping the tight pellet, containing mitochondria. This tight pellet was then resuspended in 10 mL MIBSM buffer containing 100 U of TURBO^TM^ DNase (Invitrogen) per 10 g of cells, and incubated with gentle agitation at 4°C for 20 minutes. It was then centrifuged at 10,000 x g for 15 min, 4°C before resuspending in 2 mL SEH buffer (250 mM sucrose; 1 mM EDTA; 20 mM HEPES-KOH, pH 7.4; 50 μg/mL LZD) and homogenizing gently with 5 strokes of a Dounce homogenizer. The sample was then loaded onto a chilled 15-23-32-60% sucrose gradient (prepared in 50 mM HEPES-KOH, pH 7.4; 2.5 mM EDTA; 50 μg/mL LZD) and centrifuged in a SW41 rotor at 28,500 RPM. The brown band at the 32-60% interface (containing crude mitochondria) was then collected, flash frozen in liquid nitrogen, and stored at −80°C until use.

#### Mitoribosome purification

Frozen mitochondria were thawed on ice and resuspended in 2 volumes of mitochondrial lysis buffer (25 mM HEPES-KOH, pH 7.4; 150 mM KCl; 50 mM MgOAc; 2% v/v Triton X-100; 2 mM DTT; EDTA-free protease inhibitor (Pierce); 50 μg/mL LZD) and mixed via inversion, followed by homogenization with 5 strokes of a Dounce homogenizer. The sample was then rotated on a nutator at 4°C to complete the lysis. The lysed sample was then clarified by centrifugation at 30,000 x g, 4°C, for 20 min in a TLA100.3. The supernatant was carefully removed and the pellet discarded. This centrifugation step was then repeated to ensure clarification. The clarified mitochondrial lysate was then loaded onto a sucrose cushion (1 M sucrose (34% w/v); 20 mM HEPES-KOH, pH 7.4; 100 mM KCl; 20 mM MgOAc; 1% v/v Triton X-100; 2 mM DTT; 50 μg/mL LZD) in a 2.5:1 ratio of lysate to sucrose cushion. The samples were then centrifuged at ~231,550 x g in a TLA 100.3 rotor (66,000 rpm) to pellet the mitoribosomes. The supernatant was discarded and the pellets rinsed gently several times with resuspension buffer (50 mM HEPES-KOH, pH 7.4; 250 mM KCl; 12.5 mM MgOAc; 5 mM DTT; 50 μg/mL LZD). The pellets were then resuspended in a total of 170 μL of resuspension buffer and loaded onto a 15-30% sucrose gradient (prepared in resuspension buffer using a BioComp GradientMaster^TM^) to isolate mitomonosomes. The samples were centrifuged at 19,500 rpm for 18.5 h in an SW41 rotor. The gradient was then fractionated using a BioComp Piston Gradient Fractionator^TM^ with UV260 monitoring. 200 μL fractions were collected and the fraction corresponding to the 55S peak was collected and used for cryo-EM sample preparation

### Cryo-EM sample preparation

Purified mitomonosomes were concentrated using a 100 kDa Amicon® Ultra centrifugal filter, followed by exchange with resuspension buffer (50 mM HEPES-KOH, pH 7.4; 250 mM KCl; 12.5 mM MgOAc; 5 mM DTT; 50 μg/mL LZD). The final concentration of the sample was calculated assuming that 1 A_260_ = 0.1 mg/mL^41^. Quantifoil R 1.2/1.3, copper, mesh 300 grids with 2 nm amorphous carbon layer on top were glow discharged for 30 s at 15 mA (EMS-100 Glow Discharge System, Electron Microscopy Sciences). 3 µL of a 140 nM mitoribosome sample was applied on the grid and incubated for 30 s at 100% humidity and 4°C. Variable blot times with Whatman #1 filter paper was used to control the ice thickness. Samples were vitrified with a FEI Vitrobot Mark IV (Thermo Fisher).

### Cryo-EM data processing

The dataset was collected with an FEI Talos Arctica electron microscope (200 kV, Thermo Fisher, UCSF cryo-EM core facility) using a nine-show beam image-shift approach with coma compensation. The image stacks were collected in non-super resolution mode, binned by a factor of two, motion corrected, and dose-weighted using UCSF motioncor2^42^. Dose-weighted micrographs were used to determine contrast transfer function parameters using CTFFIND 4.0 in cryoSPARC^43^. Template picker was used to pick particles corresponding to mitoribosomes. The structure 7L08^44^ was used to generate a reference map in order to generate 2D classes that were used as input for the template picker. Particles were extracted with a box size of 560 pixels (twice the largest dimension of mitoribosome in 7L08). These particles were used for 2D classification. Only classes that clearly contained ice were omitted. Homogeneous refinement was carried out in cryoSPARC using the particles corresponding to the good classes, followed by non-uniform refinement, with both global CTF and defocus refinements turned on^45^.

### Atomic model building and structure refinement

The refined and sharpened map obtained after non-uniform refinement in cryoSPARC was used for model building in Coot^46^. Restraints for linezolid were generated using eLBOW^47^ within Phenix^48^. Model refinement was performed through multiple rounds of manual model building and real space refinement in Phenix^48^. We used OPLS3e/VSGB2.1 forcefield parameters in combination with phenix.real_space_refine to improve the ligand model^49^. This approach allows a more accurate assignment of partial charges to the small molecules compared to manual restraint generation. We used standard Phenix restraints for the macromolecule while OPLS3e/VSGB2.1 forcefield was used to generate the restraints for Linezolid. We first prepared apo ribosome and Linezolid separately in Maestro prepwizard and then recombined them. The restraints for Linezolid which were generated using Maestro and the recombined complex were used as input for Phenix.real_space_refine. The protein residues and nucleotides of the 16S rRNA showed well defined geometrical parameters (Table 3). The figures were prepared using PyMol Molecular Graphics System Version 3.0.2^50^.

