## Supplementary materials for "Context-specific inhibition of mitochondrial ribosomes by phenicol and oxazolidinone antibiotics"

**This file includes:**

**Supplementary Figures:**

A. Supplementary Figure 1 (S1) – related to Figure 1
B. Supplementary Figure 2 (S2) – related to Figure 2
C. Supplementary Figure 3 (S3) – related to Figure 3
D. Supplementary Figure 4 (S4) – related to Figure 5
E. Supplementary Figure 5 (S5) – related to Figure 5
F. Supplementary Figure 6 (S6) – related to Figure 5
G. Supplementary Figure 7 (S7) – related to Figure 5

**Supplementary Tables:**

A. Supplementary Table 1: List of mRNA sequences for HiBiT reporter assay
B. Supplementary Table 2: List of tRNA sequences for HiBiT reporter assay
C. Supplementary Table 3: Cryo-EM data collection, refinement and validation statistics

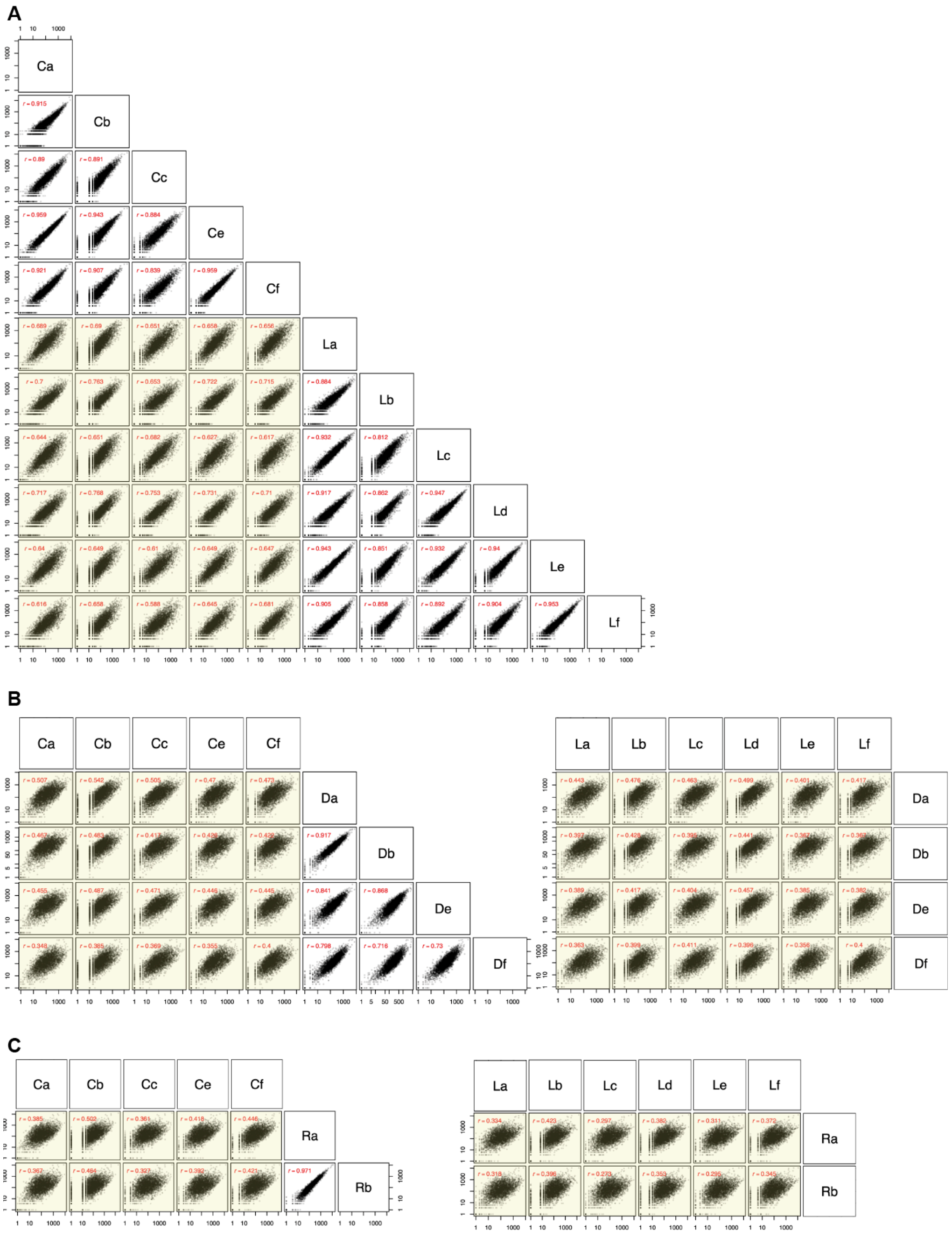


**Figure S1 – Related to Figure 1:** Replicates and oxazolidinone-treated samples are more similar than treated vs untreated or disparate treatments. (A) Pairwise correlation analysis of all chloramphenicol (CHL; Ca-Cc and Ce-Cf) vs linezolid (LZD; La, Lb-Lf) replicates. (B) Pairwise correlation analysis of all CHL and LZD replicates vs DMSO (Da, Db, De, and Df). (C) Pairwise correlation analysis of all CHL and LZD replicates vs retapamulin (RET; Ra & Rb). Pearson r is shown throughout.

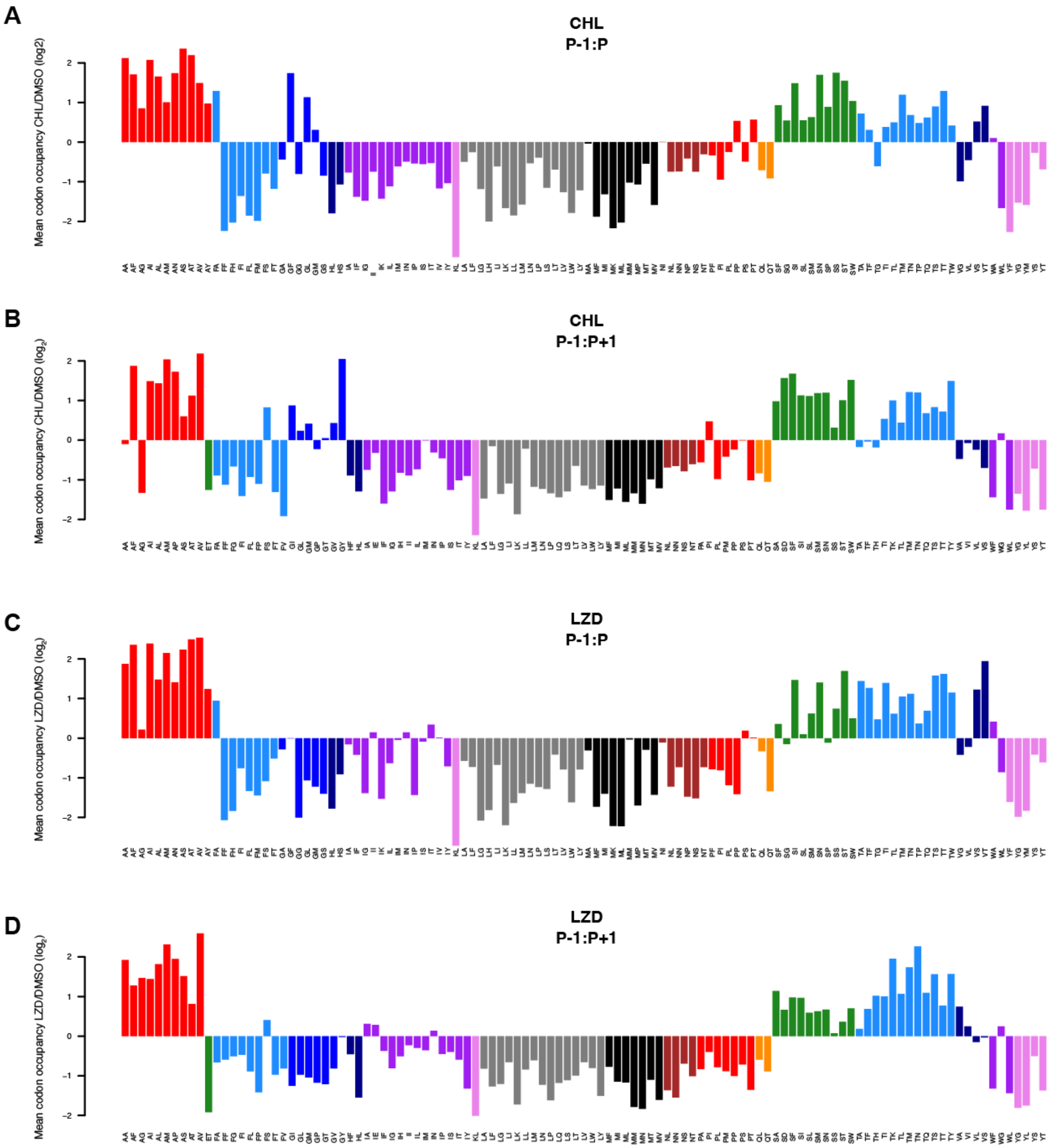


**Figure S2 – Related to Figure 2: Dicodon occupancy reveals secondary mediators of stalling by oxazolidinones.** Mean dicodon occupancies of ribosomes in CHL-treated **(A-B)** and LZD-treated **(C-D)** cells compared to DMSO are plotted for P-1 and P amino acids **(A, C)** and P-1 and P+1 amino acids **(B, D)**. A dicodon must be present at least two times in the transcriptome to be included. LZD: linezolid, CHL: chloramphenicol, DMSO: dimethyl sulfoxide

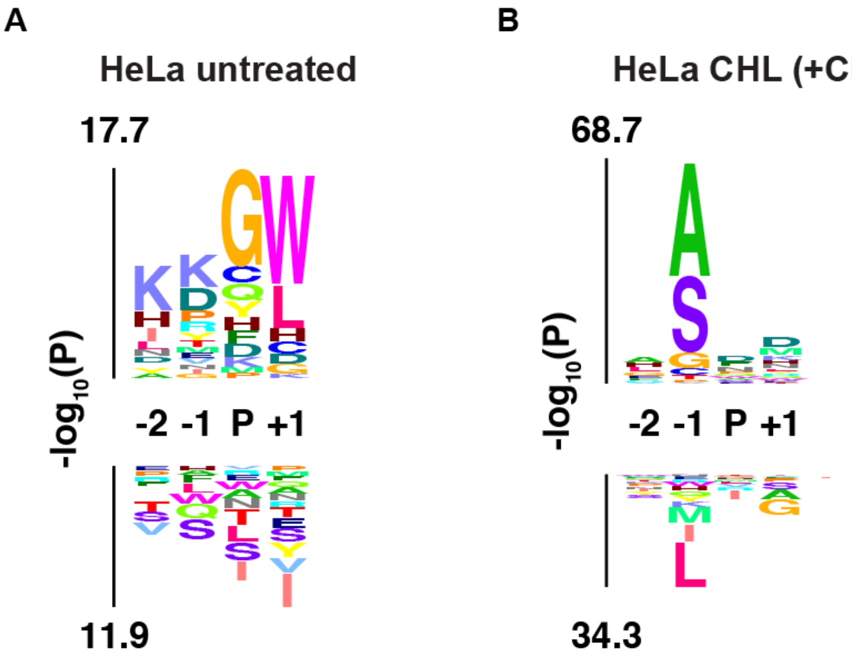


**Figure S3 – Related to Figure 3: Reanalysis of published, independent, mitoribosome profiling datasets[4] corroborate P-1 alanine stalling signature.** Mitoribosome profiling data from untreated HeLa cells (**A**) and HeLa cells treated with chloramphenicol (CHL) to halt mitoribosomal translation (100 µg/mL, 5 min) and cycloheximide (CHX) to halt cytoplasmic translation (100 µg/mL, 15 min) (**B**) were processed using the the computational pipeline developed for the present study, followed by kpLogo analysis[3] of identified stall sites. Background was taken as the average across all positions in input sequences. Y axis represents statistical significance determined by one-sided binomial tests of each residue at each position; enriched residues stack on the top, whereas depleted residues stack on the bottom.

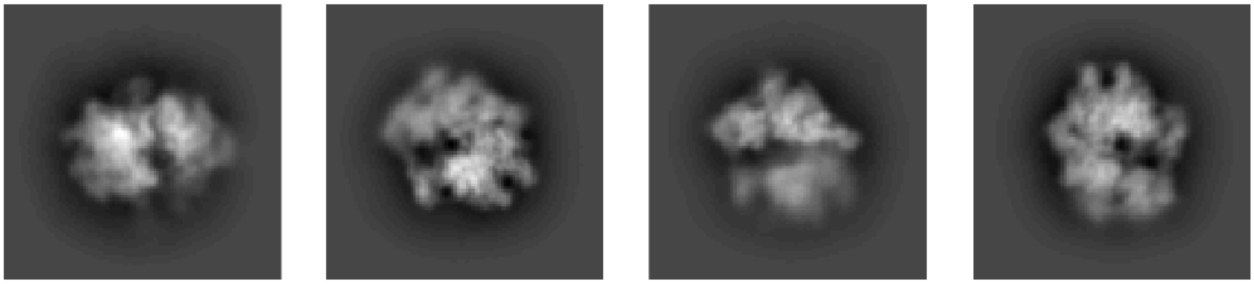

**Figure S4 – Related to Figure 5 : 2D classification showing presence of mitomonosomes.**

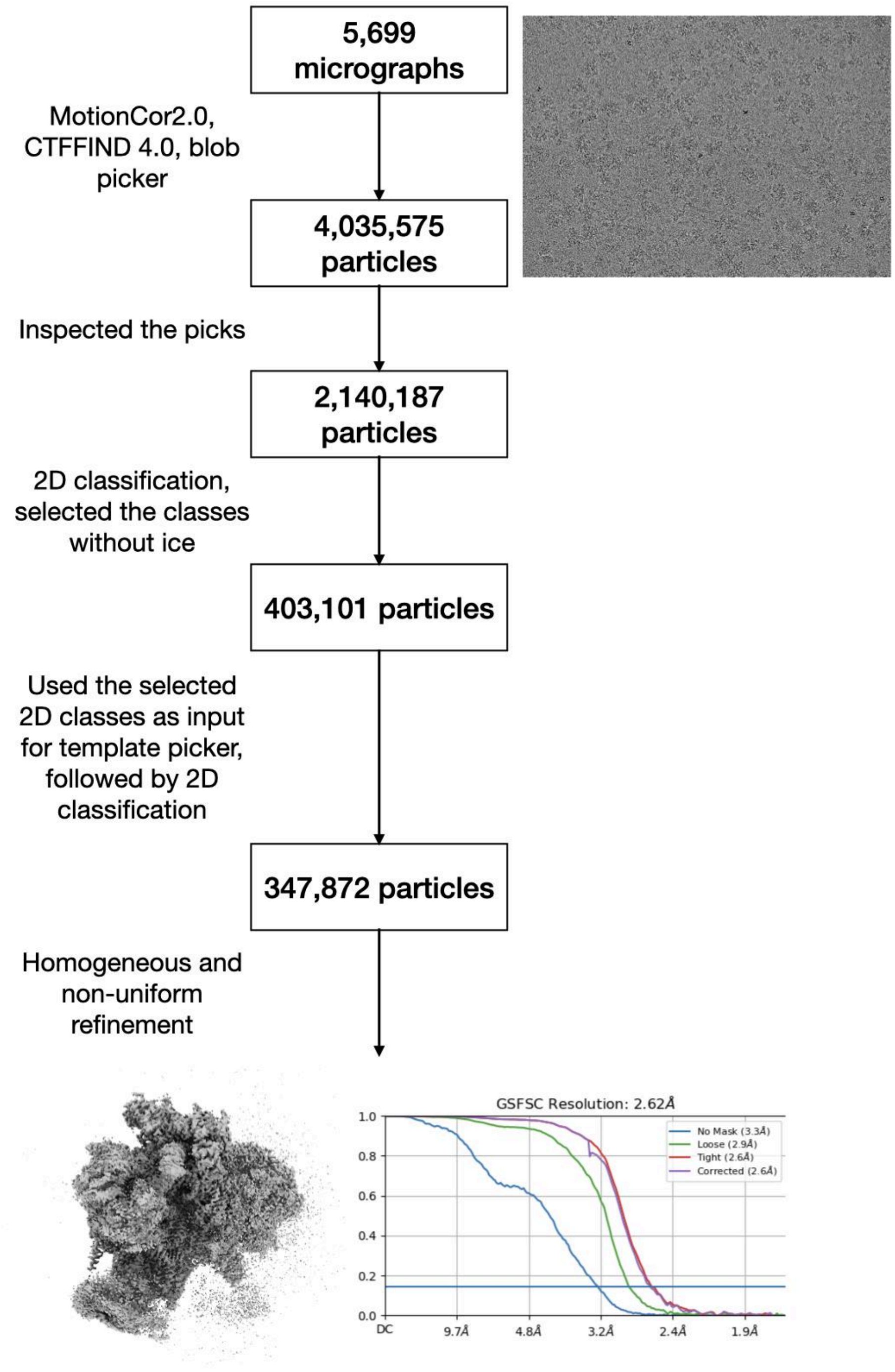


**Figure S5 – Related to Figure 5: Cryo-EM data processing workflow.**
Raw micrographs were motion corrected using MotionCor2.0 followed by CTF correction in CTFFIND 4 and curation. 2D classification was used to remove particles containing ice. The remaining good particles were used for homogeneous refinement to generate the consensus volume. The final FSC curve generated from homogeneous refinement in cryoSPARC is presented. All the steps except motion correction were performed in cryoSPARC.

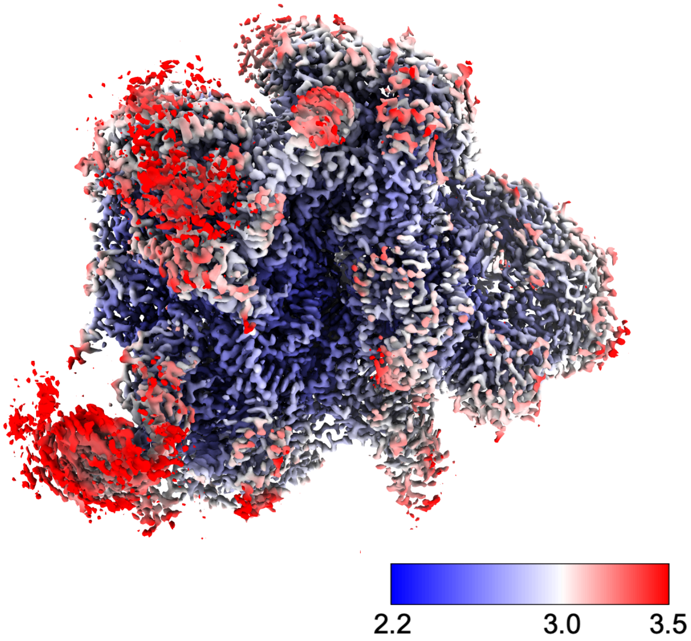


**Figure S6 – Related to Figure 5: Local resolution calculation for the final refined volume (2.2 σ).** Local resolution for the mt-LSU was calculated using the phenix.local_resolution command.

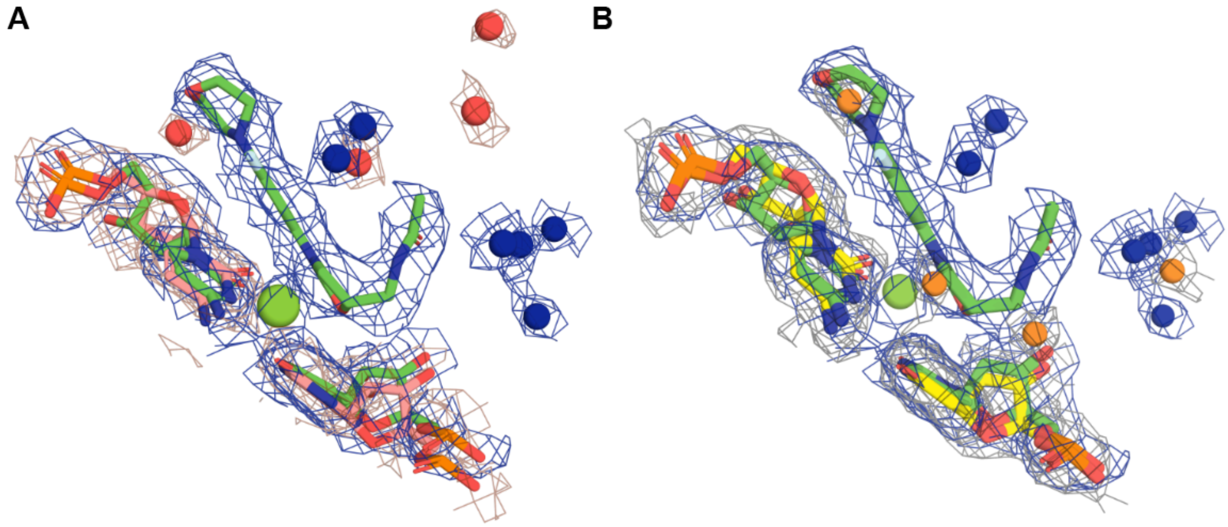


**Figure S7 – Related to Figure 5:** Overlay of present structure (Linezolid and mitoribosome nucleotides: green, waters: blue) with apo structures of the **A)** *E. coli* 50S ribosome (PDB 6PJ6; *E.coli* ribosome nucleotides: salmon, waters: red) and **B)** human mitoribosome (PDB 7QI5; mitoribosome nucleotides: yellow, waters: orange) to show waters close to ligand-interacting nucleotides in the peptidyl transfer center (PTC). Water molecules in the PTC are uniformly distributed in the apo ribosomes (apo mitoribosome and apo *E.coli* ribosome). In contrast, water molecules in the mitoribosome-LZD complex preferentially cluster around the C5 side chain of linezolid, leading to a stronger solvent density nearer the side chain. Coulomb potential density mesh is shown in blue for present structure (3.8 **σ**), brown for the apo *E. coli* 50S ribosome (2.3 **σ**) and gray for the apo human mitoribosome (3.8 **σ**).

**Supplementary Table 1: Reporter mRNAs used in mitochondrial vitro translation system**

| **mRNA** | **Sequence (5' - > 3')** |
|---|---|
| MMYALF-HiBiT (Ala_GCA) | GAUGAUGUACGCUUUAUUUGUGUCGGGUUGGCGGUUAUUCAAGAAGAUAUCGUAA$(A)_{36}$ |
| MMYYLF-HiBiT | GAUGAUGUACUACUUAUUUGUGUCGGGUUGGCGGUUAUUCAAGAAGAUAUCGUAA$(A)_{36}$ |

**Supplementary Table 2: Sequences of in vitro transcribed tRNAs (iVT tRNAs)**

| **tRNA** | **Sequence (5' - > 3')** |
|---|---|
| Yeast $tRNA^{Tyr}$ (Y) | GGGAGACCACAACGGUUUCCCUCUAUUCUCUCGGUAGC CAAGUUGGUUUAAGGCGCAAGACUGUAAAUCUUGAGAU CGGGCGUUCGACUCGCCCCCGGGAGACCA |
| Yeast $tRNA^{Ala}$ (A) | GGGCACAUGGCGCAGUUGGUAGCGCGCUUCCCUUGCAA GGAAGAGGUCAUCGGUUCGAUUCCGGUUGCGUCCACCA |
| Yeast $tRNA^{Val}$(V) | GUUCCAAUAGUGUAGCGGCUAUCACGUUGCCUUCACAC GGCAAAGGUCCCGAGUUCGAUCCUCGGUUGGAACACCA |
| Yeast $tRNA^{Gly}$(G) | GCGCAAGUGGUUUAGUGGUAAAAUCCAACGUUGCCAUC GUUGGGCCCCCGGUUCGAUUCCGGGCUUGCGCACCA |
| Yeast $tRNA^{Trp}$(W) | GAAGCGGUGGCUCAAUGGUAGAGCUUUCGACUCCAAAU CGAAGGGUUGCAGGUUCAAUUCCUGUCCGUUUCACCA |
| Yeast $tRNA^{Arg}$(R) | GCUCCUCUAGUGCAAUGGUUAGCAUGCAUUCUUCCGGU GGCUGUGAUCCGGGUUCGAGUCCCGGGAGGAGCUCCA |
| Yeast $tRNA^{Lys}$(K) | GCCUUGUUGGCGCAAUCGGUAGCGCGUAUGACUCUUAA UCAUAAGGUUAGGGGUUCGAGCCCCCUACAGGGCUCCA |
| Yeast $tRNA^{Ile}$(I) | GCUCGUGUAGCUCAGUGGUUAGAGCUUCGUGCUUAUAA CGCGACCGUCGUGGGUUCAAACCCCACCUCGAGCACCA |
| Human mito-$tRNA^{Met}$ (M) | GGUAAGGUCAGCUAAAUAAGCUAUCGGGCCCAUACCCC GAAAAUGUUGGUUAUACCCUUCCCGUACUACCA |
| Human mito-$tRNA^{Leu}$ (L) | GUUAAGAUGGCAGAGCCCGGUAAUCGCAUAAAACUUAAA ACUUUACAGUCAGAGGUUCAAUUCCUCUUCUUAACACCA |
| Human mito-$tRNA^{Phe}$ (F) | GUUUAUGUAGCUUACCUCCUCAAAGCAAUACACUGAAAA UGUUUAGACGGGCUCACAUCACCCCAUAAACACCA |
| Human mito-$tRNA^{Ser}$ (S) | GGAAAAGUCAUGGAGGCCAUGGGGUUGGCUUGAAACCA GCUUUGGGGGGUUCGAUUCCUUCCUUUUCUGCCA |

### Supplementary Table 3: Cryo-EM data collection, refinement and validation statistics

| **39S mitoribosome-Linezolid complex** | |
|---|---|
| **Data collection and processing** | |
| Facility and electron microscope | UCSF Cryo-EM core facility, Talos Arctica |
| Camera | Gatan K3 |
| Magnification | 45,000 |
| Voltage (kV) | 200 |
| Electron exposure (e-/Å2) | 21.4 |
| Defocus range (μm) | 0.5-1.2 |
| Pixel size (Å) | 0.865 |
| **Symmetry imposed** | C1 |
| **Initial particle number** | 930,137 |
| **Final particle number** | 347,872 |
| **Chains** | 116 |
| **Atoms** | 113122 (Hydrogens: 0) |
| **Residues** | Protein: 9257 Nucleotide: 1583 |
| **Water** | 4388 |
| **Ligands** | ZN: 3 |
| | ZLD: 1 |
| | MG: 103 |
| | UNK: 65 |
| **Bonds (RMSD)** | |
| Length (Å) (# > 4 σ) | 0.007 (0) |
| Angles (°) (# > 4 σ) | 0.826 (12) |
| **MolProbity score** | 2.72 |
| **Clash score** | 11.44 |
| **Ramachandran plot (%)** | |
| Outliers | 0.13 |
| Allowed | 6.4 |
| Favored | 93.47 |
| **Rama-Z (Ramachandran plot Z-score, RMSD)** | |
| whole (N = 9139) | -0.69 (0.09) |
| helix (N = 3083) | 0.96 (0.09) |
| sheet (N = 1152) | -0.38 (0.15) |
| loop (N = 4904) | -1.45 (0.09) |
| **Rotamer outliers (%)** | 3.37 |
| **Cbeta outliers (%)** | 0.03 |
| **Cis proline/general** | 1.3/0.1 |
| **Twisted proline/general** | 0.8/0.1 |

| | |
|---|---|
| **CaBLAM outliers (%)** | 3.39 |
| **ADP (B-factors)** | |
| Iso/Aniso (#) | 113122/0 |
| min/max/mean | |
| Protein | 0.00/289.60/44.38 |
| Nucleotide | 0.00/82.21/18.77 |
| Ligand | 0.00/224.64/16.25 |
| Water | 0.00/161.09/23.43 |
| **Occupancy** | |
| Mean | 1 |
| occ = 1 (%) | 99.97 |
| 0 < occ < 1 (%) | 0 |
| occ > 1 (%) | 0 |
| **Lengths (Å)** | 285.45, 259.50, 247.39 |
| **Angles (°)** | 90.00, 90.00, 90.00 |
| **Model resolution (Å)** | 2.62 |
| **FSC threshold** | 0.143 |
| **Model resolution range (Å)** | 2.4 - 3.5 |
| **Model vs. Data** | |
| CC (mask) | 0.74 |
| CC (box) | 0.67 |
| CC (peaks) | 0.64 |
| CC (volume) | 0.72 |
| Mean CC for ligands | 0.58 |
